# Distinct cochlear cell types associated with genetic susceptibility to sensory and metabolic hearing loss in older adults from the CLSA

**DOI:** 10.64898/2026.02.17.706270

**Authors:** Samah Ahmed, Kenneth I. Vaden, Judy R. Dubno, Britt I. Drögemöller

## Abstract

Hearing loss is a heterogeneous condition that can be classified into different subtypes with diverse genetic and cellular components. To investigate the cochlear cell types underlying the genetic basis of sensory and metabolic components of age-related hearing loss (ARHL), we integrated human genome-wide association study data with mouse cochlear single-cell RNA sequencing data using the single-cell disease relevance score tool. These analyses revealed that genes associated with the sensory component of ARHL in older humans were most highly expressed in the hair cells, while genes associated with metabolic component of ARHL in older humans were most highly expressed in spiral ganglion neurons. To assess whether age-related transcriptional changes might influence these patterns, we performed age-stratified analyses. In younger mice, sensory hearing loss-associated genes revealed significant heterogeneity in expression in supporting cells within the sensory epithelium. In contrast, the greatest heterogeneity in the expression of metabolic hearing loss-associated genes was observed in intermediate cells of the stria vascularis in older mice. These findings provide evidence for the role of distinct genetic and cellular risk profiles for different ARHL subtypes, suggesting that prevention and therapeutic strategies may require targeting specific cell populations at different life stages.

## Introduction

Hearing loss affects over 5% of the global population and is projected to impact nearly 2.5 billion people by 2050. Among older adults, age-related hearing loss (ARHL) is the predominant form of hearing loss.^1^ The pathophysiology of ARHL is heterogeneous, involving multiple interacting cellular and molecular mechanisms within the cochlea.^2^ Classifying ARHL into metabolic and sensory phenotypes has been proposed based on audiometric patterns.^3^ The metabolic component of ARHL primarily involves degeneration of the stria vascularis (SV) - an epithelial structure lining the lateral wall of the cochlea that generates and maintains the endocochlear potential required to power the hair cells (HCs).^4^ In contrast, the sensory component of ARHL is characterized by degeneration of or damage to HCs and associated supporting cells, which are responsible for converting mechanical vibrations into electrical signals.^5^ Because spiral ganglion neurons (SGNs) rely on both intact strial and HC function, degeneration of either of these cell types can lead to SGN impairment, impacting the transmission of auditory signals to the brain.^6^

In accordance with the differences in the biology underlying these ARHL phenotypes, recent genome-wide association studies (GWAS) performed by our group identified distinct genetic variants and genes associated with metabolic and sensory components of ARHL.^7^ To better understand each of the hearing loss phenotypes at the genetic level and inform treatment approaches, it is essential to determine the specific cell types that express the ARHL subtype-associated genes. In one straightforward example, this approach has been applied to the treatment of Mendelian deafness, where identification of *OTOF*, a gene that is uniquely expressed in HCs, paved the way for successful targeted gene therapy in some patients.^8^ Unfortunately, our understanding of the genes and cell types underlying ARHL remains limited, which continues to limit the development of precise and effective treatments.

Single-cell RNA sequencing (scRNA-seq) data provide high-resolution gene expression profiles across diverse cochlear cell populations, including rare but critical inner and outer HCs.^9^ Integrating scRNA-seq mouse datasets with GWAS findings from humans has become an important strategy to identify cochlear cell types potentially involved in ARHL, based on the principle that cell types with elevated expression of top GWAS-mapped genes may contribute to the underlying pathophysiology. Despite the promise of these strategies, previous studies using human GWAS datasets have produced contradictory results. Kalra *et al*. reported that cells of the sensory epithelium (e.g., HCs and supporting cells) are the primary cell types implicated in ARHL.^10^ In contrast, Trpchevska *et al*. found that lateral wall cells (e.g., cells of the SV) are mainly involved in ARHL, with no evidence for the enrichment of sensory epithelia.^11^ In an effort to resolve this discrepancy, Hertzano *et al*. performed the same analyses on mouse cochlear samples including all major cell types.^12^ Similar to Kalra *et al*., they found that sensory epithelial cells are more likely to be involved in the genetic basis of ARHL, even though it is known that the SV plays an important role in this condition.^13,14^ These discrepancies may, in part, arise from the use of measures of self-reported hearing difficulty rather than pure-tone audiometry, which do not allow determination of sensory and metabolic components of ARHL.

To provide more detailed insight on the cochlear cell types underlying the genetic basis of ARHL, we integrated human GWAS findings derived from accurate audiometric phenotyping of metabolic and sensory estimates of ARHL with murine inner ear scRNA-seq expression profiles. This approach enabled us to investigate the contribution of specific cell types to sensory and metabolic ARHL to unravel the cellular bases of these two distinct ARHL phenotypes.

## Methods

### GWAS and scRNA-seq data sources

scRNA-seq expression profiles were obtained from a previously published study by Sun *et al*., which generated a comprehensive atlas of the aging mouse cochlea.^15^ The data from this study is publicly available through the Genome Sequence Archive (GSA:CRA004814). In brief, cochlear samples were collected from C57BL/6J mice at five time points: 1-2 months (young adult), 5 months (middle-aged), and 12-15 months (aged), with equal representation of males and females. In total, 45,972 single-cell transcriptomes were generated from 27 cochlear cell types. Preprocessing steps in the original study included stringent quality control to retain high-quality cells and perform data normalization. To enable meaningful biological comparisons, data were integrated across samples. Dimensionality reduction and clustering were then performed to identify cell types based on classic marker genes. The preprocessed and cell-type annotated scRNA-seq object was retrieved from the gEAR database (https://umgear.org/) and used directly as input for downstream analyses.^16^

GWAS summary statistics for sensory and metabolic ARHL phenotypes were obtained from our previous study by Ahmed *et al*.^7^ In short, the sensory and metabolic components of ARHL were calculated from audiograms of participants in the Canadian Longitudinal Study on Aging (CLSA) by fitting each audiogram to previously developed profiles by Vaden *et al*.^3,17,18^ GWAS was then performed using a linear regression model. The resulting GWAS summary statistics were used for gene-based association testing with MAGMA, in which genetic variants were mapped to their corresponding genes and aggregated into gene-level Z-scores and *P*-values, as previously described.^7^

### Mapping human genes to mouse orthologs

Given that the scRNA-seq data used in this study were derived from murine models, GWAS-prioritized human genes identified through MAGMA analysis were mapped to their corresponding mouse orthologs (*Mus musculus* C57BL/6J) using Ensembl BioMart (https://useast.ensembl.org/biomart/martview/). To further assess cross-species similarity, we retrieved the transcript and protein reference FASTA sequences for significant MAGMA genes from the NCBI database for humans and mice (https://www.ncbi.nlm.nih.gov/). Then, genomic sequences of human-mouse ortholog pairs were aligned using BLAST to determine sequence identity (similarity) and coverage (the percentage of the human sequence aligning to the corresponding mouse sequence).^19^

### GWAS and scRNA-seq data integration

We integrated the murine scRNA-seq expression data with our human GWAS results using the single-cell Disease Relevance Score (scDRS) method.^20^ scDRS estimates whether individual cells exhibit excess expression of trait-associated genes by computing scores based on the cumulative expression of the prioritized genes across the genome. The method generates two key metrics: (1) association scores that measure the strength of enrichment of the disease score for each cell type, and (2) heterogeneity scores that assess variability in the disease score across individual cells within each cell type cluster.

For association score analyses in scDRS, we ranked genes previously identified by Ahmed *et al*. based on their association statistics and selected the top 1000 genes from the MAGMA gene-based results, consistent with the default threshold recommended by scDRS.^7^ The scRNA-seq dataset of the entire mouse cochlea was used for these analyses to maximize statistical power. Additionally, we evaluated smaller and more specific MAGMA-derived gene sets for these association score analyses as follows: (1) top 20 genes scDRS and (2) top 100 genes, corresponding to a gene-set that is 10-fold smaller than the scDRS default.

To assess cell-type-specific changes across time, we performed scDRS heterogeneity analysis using the top 1000 MAGMA-derived genes across three murine scRNA-seq age-stratified groups: 1-2 months (young adult), 5 months (middle-aged), and 12-15 months (aged). Multiple testing correction for 27 cells was performed for all analyses using the Benjamini-Hochberg false discovery rate (FDR) method, with FDR *P* < 0.05 considered statistically significant.

## Results

### Identification of cross-species gene mapping and sequence conservation

Of the top 1000 MAGMA genes, 971 sensory ARHL genes and 964 metabolic ARHL genes had Ensembl annotations. Among these, 106 sensory and 90 metabolic ARHL genes lacked mouse orthologs, leaving 865 sensory 874 and metabolic ARHL genes for downstream analysis (**Supplementary Tables 1 and 2**). Sequence comparison between human and mouse orthologs for the significant genes identified by MAGMA^7^ (*P*<2.57×10^-6^) demonstrated strong homology at the transcript and protein levels, with sequence identity exceeding 70% (**Supplementary Table 3**). This indicates that the molecular architecture of these genes is largely preserved across species. The only exception to this was *KLHDC7B*, the top gene associated with sensory ARHL, which although showing high sequence similarity at the transcript level (80.1%), exhibited a lower degree of protein sequence similarity (51.9%) relative to the other genes, suggesting potential species-specific divergence at the protein level. Nonetheless, the amino acid affected by the lead GWAS variant in this gene (rs36062310; p.Val1145Met; *P*=2.37×10^-12^) remains well conserved. Overall, these results support the validity of using mouse single-cell transcriptomic data to explore the expression patterns of human genes associated with ARHL.

### Cell type prioritization in metabolic and sensory ARHL

To identify cell types that contribute to the genetic bases of sensory and metabolic ARHL, we integrated MAGMA gene-level association results with mouse cochlear scRNA-seq data using scDRS. Among the top genes, 155 sensory ARHL genes (including *KLHDC7B* and *MYO15A*) and 152 metabolic ARHL genes were filtered out due to low expression in the scRNA-seq dataset, retaining 710 sensory and 722 metabolic genes for scDRS analysis (**Supplementary Tables 1 and 2**). These analysis of the remaining genes revealed that the cell types that were uncovered as most important for the each of the ARHL subtypes were distinct. For sensory ARHL, HCs consistently showed the strongest association signals across gene-set sizes, with the gene-set containing the top 100 genes showing the most significant results (*P* = 0.003 FDR adjusted *P* = 0.081) (**Figure 1 A-C, Supplementary Table 4**). In contrast, for metabolic ARHL, SGNs were uncovered as the most important cell type across gene-set sizes, with the gene-set containing the top 20 genes showing the most significant results (*P* = 0.003; FDR adjusted *P* = 0.081) (**Figure 1 D-F, Supplementary Table 5**).

**Figure 1:**
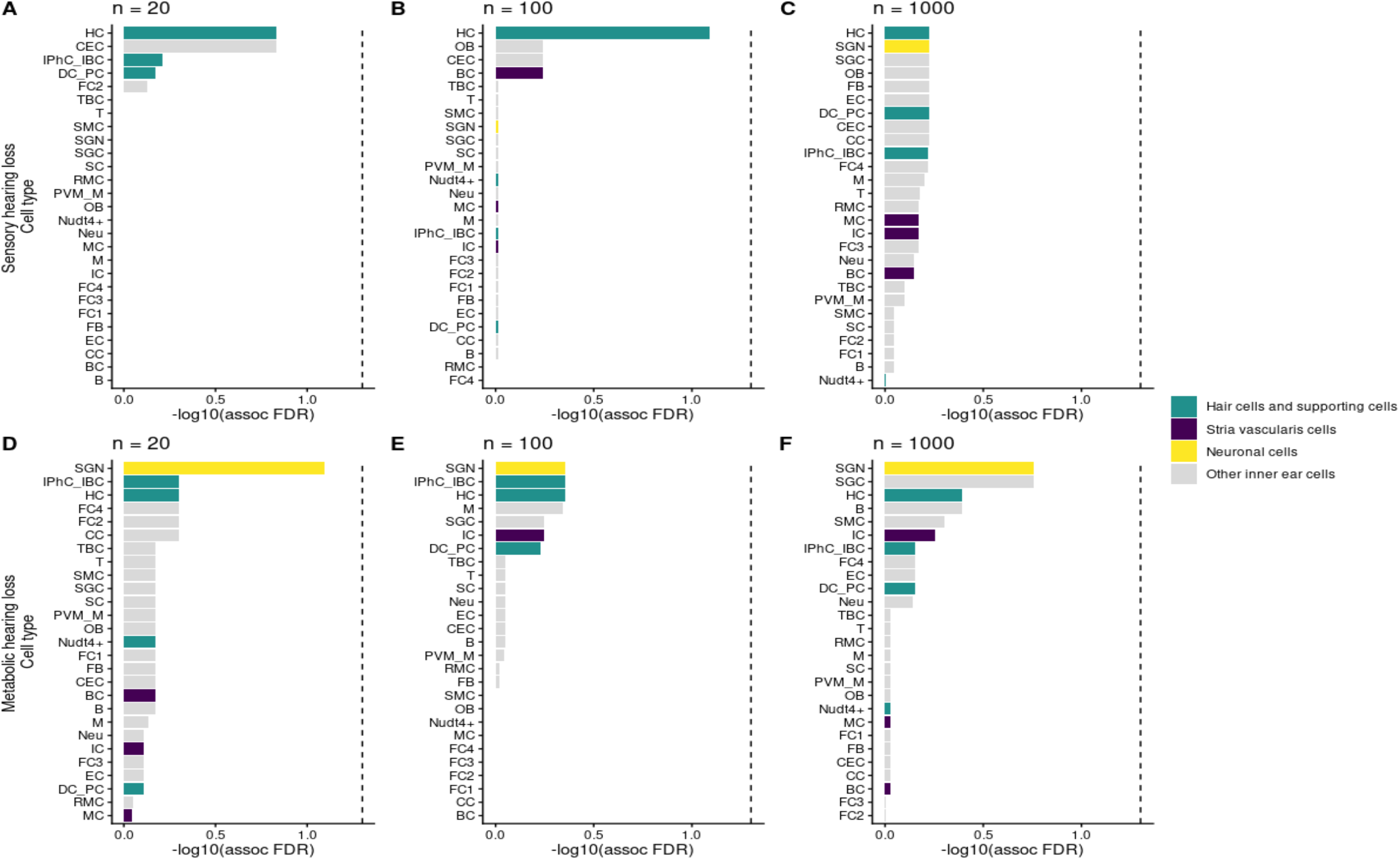
Cell type-specific association of sensory (A-C) and metabolic (D-F) hearing loss for different gene-set sizes. The dashed line indicates the FDR-corrected significance threshold -log10(0.05). **\*Cell types: HC**: Hair cells, **DC_PC**: Dieter cells and pillar cells, **IPhC_IBC**: Inner phalengeal cells and inner border cells, **TBC**: Tympanic border cells, **Nudt4+**: Nudt4 pillar cells, **EC**: Epithelial cells, **SGN**: Spiral ganglion neurons, **SGC**: Satellite glial cells, **SC**: Schwann cells, **CC**: Chondrocytes, **OB**: Osteoblasts, **RMC**: Reissner’s membrane cells, **IC:** Intermediate cells, **MC**: Marginal cells, **BC**: Basal cells, **CEC**: Capillary endothelial cells, **PVM_M**: Perivascular resident macrophage-like melanocytes, **FB**: Fibroblasts, FC1: Fibrocytes1, **FC2**: Fibrocytes2, **FC3:** Fibrocytes3, **FC4**: Fibrocytes4, **SMC**: Smooth muscle cells, M: Macrophages, **T**: T cells, **B**: B cells, Neu: Granulocytes, **Neu**: Neutrophils

Heterogeneity analyses of the top 1000 MAGMA-derived genes also revealed distinct cell-type-specific enrichment patterns for the two hearing-loss phenotypes across age groups. For sensory ARHL, Nudt4+ supporting cells - a subtype of pillar/supporting cells within the sensory epithelium - showed significant heterogeneity in gene expression in the youngest group of 1-2-month-old mice (*P* = 0.001; FDR adjusted *P* = 0.027; **Figure 2 A-C, Supplementary Table 6**). This indicates that heterogeneity in the expression of sensory ARHL genes occurs in the sensory-epithelial cells early in life. In contrast, although not statistically significant (*P* = 0.003; FDR adjusted *P* = 0.081), metabolic ARHL genes exhibited heterogeneity in the expression in the intermediate cells (ICs) of the SV in the oldest group (12-15-month-old mice). This highlights the increasing contribution of strial dysfunction with age for metabolic ARHL (**Figure 2 D-F, Supplementary Table 7**).

**Figure 2:**
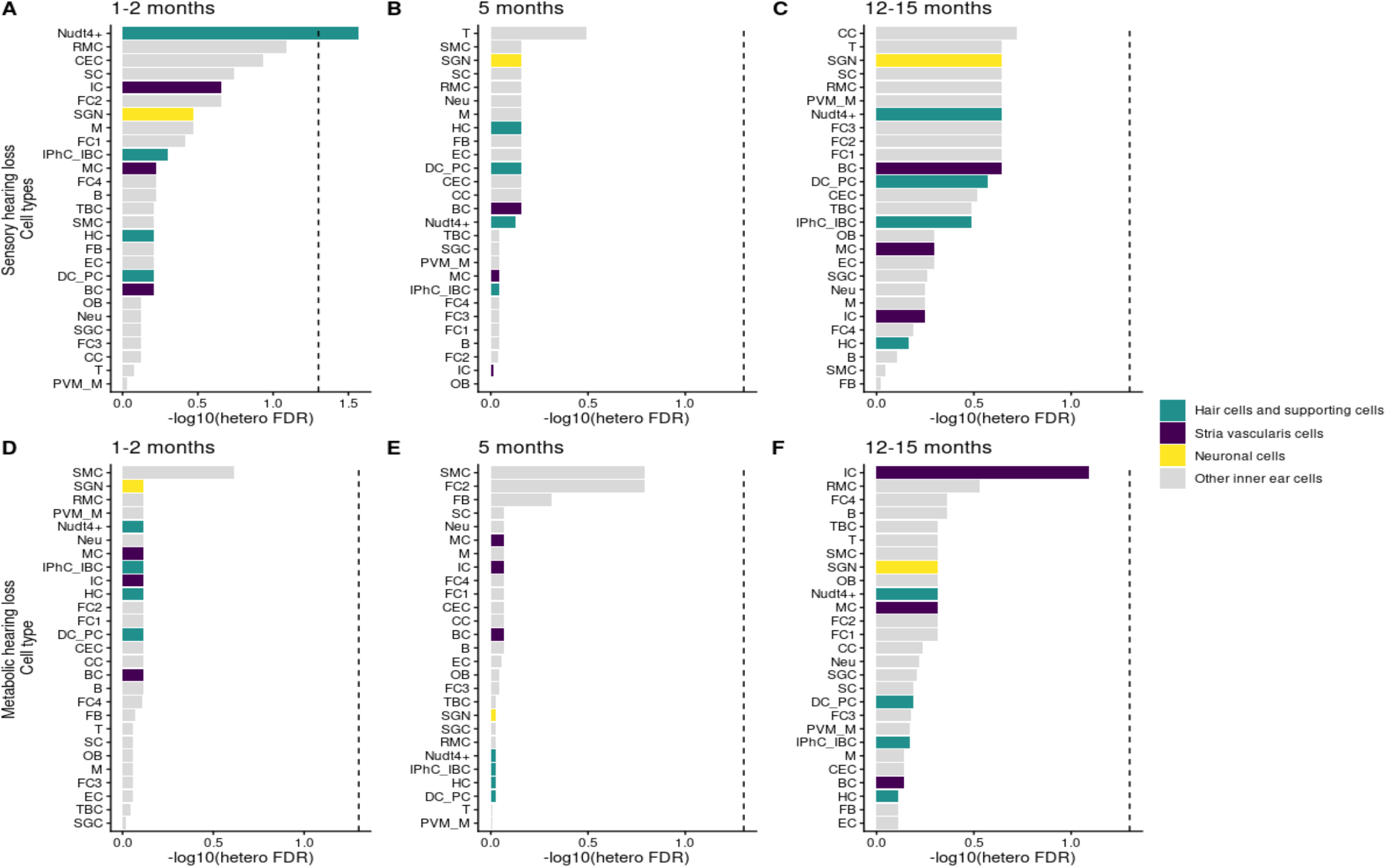
Cell type-specific heterogeneity of the top 1,000 genes for sensory (A-C) and metabolic (D-F) phenotypes. The dashed line indicates the FDR-corrected significance threshold -log10(*0*.*05*). **\*Cell types: HC**: Hair cells, **DC_PC**: Dieter cells and pillar cells, **IPhC_IBC**: Inner phalengeal cells and inner border cells, **TBC**: Tympanic border cells, **Nudt4+**: Nudt4 pillar cells, **EC**: Epithelial cells, **SGN**: Spiral ganglion neurons, **SGC**: Satellite glial cells, **SC**: Schwann cells, **CC**: Chondrocytes, **OB**: Osteoblasts, **RMC**: Reissner’s membrane cells, **IC:** Intermediate cells, **MC**: Marginal cells, **BC**: Basal cells, **CEC**: Capillary endothelial cells, **PVM_M**: Perivascular resident macrophage-like melanocytes, **FB**: Fibroblasts, FC1: Fibrocytes1, **FC2**: Fibrocytes2, **FC3:** Fibrocytes3, **FC4**: Fibrocytes4, **SMC**: Smooth muscle cells, M: Macrophages, **T**: T cells, **B**: B cells, Neu: Granulocytes, **Neu**: Neutrophils

## Discussion

This study revealed distinct cell types contribute to the genetic basis of sensory and metabolic ARHL. By integrating mouse scRNA-seq data, including all cochlear cell types, with GWAS-prioritized genes based on audiogram-derived sensory and metabolic estimates, we conducted a comprehensive assessment of cell-type-specific enrichment in ARHL subtypes. This approach has played an important role in providing additional context for the conflicting results using similar techniques reported in recent publications.^8-10^ The results obtained from these detailed scDRS analyses and GWAS results from a large dataset of older adults indicate that HCs are the most important cell type for the genetic basis of sensory ARHL. In contrast, we found that SGNs were the cell type most strongly associated with metabolic ARHL (**Figure 1**).

The involvement of HCs in the sensory phenotype aligns with their essential role in mechanoelectrical transduction and their well-known vulnerability to noise and ototoxins.^5^ Because degeneration of outer and inner HCs is a primary driver of the sensory component of ARHL, these cells are expected to show the strongest genetic signal in this phenotype. The association between cells in the SGN and metabolic ARHL is also biologically plausible. Metabolic ARHL is primarily driven by strial atrophy and reduced endocochlear potential,^4,21^ placing chronic stress on SGNs, leading to secondary neuronal degeneration.^22^ The involvement of the SGNs that was uncovered by this study therefore likely reflects the downstream effects of metabolic decline in strial cells.

Additional analyses using scDRS to investigate the heterogeneity of sensory and metabolic ARHL-associated gene expression also revealed phenotype-specific patterns consistent with the diverse genetic architecture of the two phenotypes. These patterns provide important insight into the temporal dynamics of ARHL risk. The early involvement of sensory-epithelial subtypes in the sensory phenotype suggests that genetic susceptibility may exert its effects relatively early in life, with sensory ARHL-associated genes expressed more highly in certain populations of sensory epithelial cells. This heterogeneity in expression may reflect the spatial gradient of degeneration of HCs observed in sensory hearing loss, where cells in the high-frequency or basal regions of the cochlea deteriorate first.^3,15^

In contrast, the progressive emergence of strial involvement in the metabolic phenotype is consistent with the known vulnerability of the SV to age-related metabolic decline.^3^ This age-dependent shift suggests that while metabolic genetic risk may initially manifest in sensory and neuronal cells, aging introduces a heterogeneous effect on specific strial cell populations (**Figure 2**). This pattern is also consistent with cell deterioration occurring gradually over time, where changes in gene expression arising in one cell may trigger a cascade of downstream effects that impact adjacent cells.^23^ Together, these observations highlight that the cellular involvement of genetic risk for ARHL shifts across the lifespan. They also suggest that preventive and therapeutic strategies for sensory versus metabolic ARHL may need to target distinct cell populations at different life stages.

While our study provides valuable insights into the cell-type enrichment of different phenotypes of ARHL; several limitations should be noted. First, although cross-species sequence analysis demonstrated high conservation for most genes, reliance on mouse scRNA-seq data remains an inherent limitation of this study, with differences in cochlear anatomy, gene regulation, and developmental timing potentially impacting results.^24,25^ In addition, the C57BL/6J mouse strain and related mouse models exhibit accelerated ARHL as early as 3-6 months of age, which differs from the typical temporal pattern observed in human.^26^ Therefore, future investigations would benefit from the inclusion of other mouse strains, which shown patterns of ARHL more consistent with humans. Second, several key ARHL associated genes (e.g., *KLHDC7B* and *MYO15A*) exhibited minimal or no expression in the mouse scRNA-seq dataset used in this study, which makes it difficult to assess their impact on cochlear cell function. Third, the small sample sizes of the available cochlear single-cell datasets likely limited statistical power, contributing to the lack of significant associations after multiple-testing correction.

## Conclusion

Our results identified distinct cochlear cell types involved in sensory and metabolic ARHL, with HCs most strongly associated with the sensory phenotype and SGNs with the metabolic phenotype. We also observe age-dependent heterogeneity in gene-expression-based associations, particularly within HCs for sensory and ICs for metabolic ARHL. Future work using spatial transcriptomics will be important for confirming and extending these findings.

## Acknowledgements and funding

This research was made possible using the data/biospecimens collected by the Canadian Longitudinal Study on Aging (CLSA). Funding for the CLSA is provided by the Government of Canada through the Canadian Institutes of Health Research (CIHR) under grant reference: LSA 94473 and the Canada Foundation for Innovation, as well as the following provinces, Newfoundland, Nova Scotia, Quebec, Ontario, Manitoba, Alberta, and British Columbia. This research has been conducted using the CLSA Baseline Comprehensive Dataset Version 6.0 and Genome-wide Genetic Data Version 3.0 under Application Number 2104035. The CLSA is led by Drs. Parminder Raina, Christina Wolfson and Susan Kirkland. The opinions expressed in this manuscript are the author’s own and do not reflect the views of the Canadian Longitudinal Study on Aging. The authors gratefully acknowledge the time and commitment of the CLSA participants, without whom this research would not be possible.

This work was funded through a Natural Sciences and Engineering Research Council of Canada Discovery Grant (B.I.D.) (RGPIN-2022-04500), Canadian Institutes of Health Research Catalyst Grant (B.I.D) (AFF-187272) and (in part) by the National Institutes of Health/National Institute on Deafness and Other Communication Disorders Clinical Research Center (P50 DC 000422) awarded to the Medical University of South Carolina and by the South Carolina Clinical and Translational Research (SCTR) Institute, with an academic home at the Medical University of South Carolina, NIH/NCATS Grant number UL1 TR001450 (J.R.D. and K.V.). Portions of this investigation were conducted in a facility constructed with support from Research Facilities Improvement Program Grant Number C06 RR14516 from the NIH/NCRR (J.R.D. and K.V.). B.I.D. is supported by a CIHR Tier 2 Canada Research Chair in *Pharmacogenomics and Precision Medicine* (CRC-2019-00040/CRC-2023-00351). S.A. was supported through a Research Manitoba Studentship at the University of Manitoba.

## Author contributions

S.A. performed the discovery analyses and wrote the manuscript. K.I.V. and J.R.D. developed the phenotyping approach and provided clinical expertise. B.I.D. conceived of and supervised the project. All authors contributed to editing the manuscript.

## Data availability

Data are available from the Canadian Longitudinal Study on Aging (www.clsa-elcv.ca) for researchers who meet the criteria for access to de-identified CLSA data in addition to imputed STRs. Scripts that were used in this study are available on https://github.com/Drogemoller-Lab/.

